# Emergence and suppressive function of Tr1 cells in glomerulonephritis

**DOI:** 10.1101/2023.02.07.527417

**Authors:** Shiwa Soukou-Wargalla, Christoph Kilian, Lis Velasquez, Andres Machicote, Franziska Bertram, Friederike Stumme, Tanja Bedke, Anastasios Giannou, Jan Kempski, Morsal Sabihi, Ning Song, Hans-Joachim Paust, Alina Borchers, Laura Garcia Perez, Penelope Pelczar, Beibei Liu, Can Ergen, Babett Steglich, Franziska Muscate, Tobias B. Huber, Ulf Panzer, Nicola Gagliani, Christian F. Krebs, Samuel Huber

**Affiliations:** I. Department of Medicine, University Medical Center Hamburg-Eppendorf, Hamburg, Hamburg, Germany; Hamburg Center for Translational Immunology (HCTI), University Medical Center Hamburg-Eppendorf, Hamburg, Germany; Department for General, Visceral and Thoracic Surgery, University Medical Center Hamburg-Eppendorf, Hamburg, Germany; III. Department of Medicine, University Medical Center Hamburg-Eppendorf, Hamburg, Germany

## Abstract

T regulatory type 1 (Tr1) cells, which are defined by their regulatory function, lack of Foxp3, high expression of IL-10, CD49b, and LAG3, are known to be able to suppress Th1 and Th17 in the intestine. Th1 and Th17 cells are also the main drivers of crescentic glomerulonephritis, the most severe form of renal autoimmune disease. However, whether Tr1 cells emerge in renal inflammation and moreover, whether they exhibit regulatory function during glomerulonephritis has not been thoroughly investigated yet. To address these questions, we used a mouse model of experimental crescentic glomerulonephritis and double Foxp3^mRFP^ IL-10^eGFP^ reporter mice. We found that Foxp3^neg^ IL-10-producing CD4^+^ T cells infiltrate the kidneys during glomerulonephritis progression. Using single-cell RNA- sequencing, we could show that these cells express the core transcriptional factors characteristic of Tr1 cells. In line with this, Tr1 cells showed a strong suppressive activity *ex vivo* and were protective in experimental crescentic glomerulonephritis *in vivo*. Finally, we could also identify Tr1 cells in the kidneys of patients with anti-neutrophil cytoplasmic autoantibody (ANCA)-associated glomerulonephritis and define their transcriptional profile. Tr1 cells are currently used in several immune-mediated inflammatory diseases, e.g. as T- cell therapy. Thus, our study provides proof of concept for Tr1 cell-based therapies in experimental glomerulonephritis.

## Introduction

Immune-mediated inflammatory diseases (IMIDs) are characterized by an expansion of pro-inflammatory CD4^+^ T-cell subsets, such as Th1 and Th17 cells. Accordingly, inflammatory diseases in the kidney, such as anti-neutrophil cytoplasmatic antibody (ANCA)-associated glomerulonephritis (GN), lupus nephritis, and anti-glomerular basement membrane (anti-GBM) glomerulonephritis are characterized by an increase of effector CD4^+^ T cells, especially of Th1 and Th17 cells (1–5). Thus, one key aim is to understand how these cells can be therapeutically controlled. The immune system has several mechanisms by which effector CD4^+^ T cells can be controlled in order to maintain and reestablish homeostasis. A first mechanism to control effector CD4^+^ T cells is exerted via regulatory T cells. Foxp3^+^ regulatory T cells (Foxp3^+^ Treg) are one anti-inflammatory T cell subset, which has been extensively studied in the context of GN (6–9). It has been shown that Foxp3^+^ Treg via the production of IL-10 control Th17 cells in the kidney and the intestine (10, 11). There is another anti-inflammatory T-cell subset, which is also known to produce high amounts of IL-10, namely regulatory type 1 T cells, Tr1 cells. Tr1 cells are CD4^+^ T cells that are defined by the absence of Foxp3, expression of IL-10, and regulatory function (12, 13). Interestingly, the suppressive activity of Tr1 cells also depends on functional IL-10 receptor signaling (11, 14, 15). In fact, when Tr1 cells cannot respond to IL-10, they lose the capacity to maintain their own IL-10 production, consequently their suppressive activity declines (14). Mouse Tr1 cells differentiation can be induced *in vitro* by the addition of the cytokine IL-27 (16). Tr1 cells are identified via a combination of criteria: high IL-10 expression, lack of Foxp3, co- expression of the integrin alpha2 subunit (CD49b), and the lymphocyte-activation gene 3 (LAG-3) (13, 17). Additionally, there are more co-inhibitory receptors that are known to be expressed by Tr1 cells, such as T cell immunoglobulin and ITIM domain (TIGIT), the transmembrane protein TIM-3 (TIM3), programmed cell death protein 1 (PD-1) and C-C chemokine receptor type 5 (CCR5).

Recently, the presence of Tr1 cells has been identified in the kidney of patients with IgA vasculitis (18). IL-10^+^ CD4^+^ T cells have also been shown to expand and mediate a regulatory function in patients with lupus nephritis after methylprednisolone therapy (19). Similarly, Foxp3^+^ and Tr1-like cells expand after nasal myeloperoxidase (MPO) peptide-tolerization therapy in a murine MPO-ANCA GN model and reduce disease severity (20). Thus, Tr1 cells seem to be present in different forms of renal autoimmune disease and might mediate tolerogenic mechanisms. However, whether these cells are *bona fide* Tr1 cells is not clear since the methods for identifying them mainly relied on the expression of IL-10 and lack of Foxp3 co-expression via flow cytometry or CD49b and LAG-3 co-stain by immunofluorescence. Furthermore, if these cells are able to suppress renal inflammation is unclear. Finally, the origin of Tr1 cells in the kidney is unknown. Indeed, Th17 cells can convert into regulatory T cells, referred to as Tr1^exTh17^ cells in the intestine (21).

Therefore, we aimed here to study the emergence and function of Tr1 cells in the context of glomerulonephritis. By doing so, we found that Tr1 cells emerge in the kidney, but only a small fraction of these cells are derived from Th17 cells. Moreover, functional *in vitro* as well as *in vivo* analysis confirmed that Tr1 cells have regulatory activity in the kidney in a mouse GN model. Finally, we showed the existence of Tr1 cells in human kidney biopsies. Thus, this study forms the basis to target Tr1 cells in patients with glomerulonephritis.

## Materials & Methods

### Animals

Mice were kept under specific pathogen-free conditions in the animal research facility of the University Medical Center Hamburg-Eppendorf (UKE). Food and water were provided *ad libitum*. *Rag1^-/-^* were obtained from the Jackson Laboratory. Foxp3^mRFP^, IL17a^eGFP^, IL17A^Katushka^, IL10^eGFP^ reporter mice, and IL17A^Cre^, Rosa26^YFP^ are described elsewhere (13, 21–24). Age and sex-matched littermates between 8-12 weeks were used. All animals were cared for in accordance with the institutional review board ‘Behörde für Soziales, Familie, Gesundheit und Verbraucherschutz’ (Hamburg, Germany) listed under the animal protocol number 17/2012.

### Induction of experimental crescentic glomerulonephritis and functional studies

For the induction of the experimental crescentic glomerulonephritis (nephrotoxic nephritis, NTN), 8-12 weeks- old male mice were injected intraperitoneally with an anti-serum raised against the glomerular basement membrane.

### Anti-CD3 specific antibody mouse model

Mice were injected intraperitoneally with 15 µg anti-CD3 specific antibody (Clone 2C11) dissolved in 100 µl PBS. Injections were performed on days 8 and 10 after induction of glomerulonephritis. Mice were sacrificed four hours after the second injection was applied.

### T cell-differentiation and *in vivo* transfer model

Firstly, isolated spleens and lymph nodes were smashed through a 100 µm cell strainer. Total cells were pelleted. For the transfer of CD4^+^ T cells, first CD4^+^ T cells were isolated via positive selection using Magnetic activated cell sorting (MACS) (Miltenyi Biotec, Bergisch Gladbach, Germany). APCs were isolated with the same system via CD4 and CD3 negative selection. To avoid proliferation during *in vitro* culture, APCs were irradiated with 30 Gy.

For Th17 cellproliferation, plates were coated with PBS containing 2 µg/ml anti-CD3 specific antibody. Plates were incubated overnight at 4°C or for 3 hours at 37°C. 1x10^6^ naïve CD4^+^ T cells/ml isolated from reporter mice were cultured in full Click’s medium that was supplemented with 10 % FBS, 1 % l- glutamine, 1 % penicillin/streptomycin, and 1:1000 β- Mercaptoethanol. 4x10^6^ irradiated APCs/ml, anti-CD3 specific antibody (3 µg/ml), anti-CD28 (2 µg/ml), TGF-β (0,5 ng/ml), IL-6 (10 ng/ml), IL-23 (20 ng/ml), anti-IFN- γ (10 µg/ml), and anti-IL-4 (10 µg/ml) were added to the full Click’s medium. Cells were cultured for 5 days at 37°C – 5 % CO_2_.

For Tr1 cell proliferation, 24 well plates were coated with PBS containing 2 µg/ml anti-CD3 specific antibody. Plates were incubated overnight at 4°C or for 3 hours at 37°C. Prior to culture, the coating solution was removed completely. After depletion of CD25^+^ cells, 1x10^6^ CD4^+^ T cells/ml from reporter mice were resuspended in full Click’s medium that contained anti-CD28 (2 µg/ml) and IL-27 (30 ng/ml). Cells were cultured for 5 days at 37°C 5 % CO_2_. The expression of Foxp3^mRFP^ and IL-10^eGFP^ was determined using flow cytometry.

Fluorescence-activated cell sorting (FACS-sorting) was performed on a BD FACS Aria Illu or AriaFusion. Th17 cells were FACS-sorted by negative expression of Foxp3^mRFP^ and positive expression for IL-17A^eGFP^. Foxp3^-^ Tr1 cells were sorted for high expression of IL- 10^eGFP^. Foxp3^+^ Tregs cells were isolated from spleens of reporter mice and sorted according to Foxp3^mRFP^ expression. For the *in vivo* transfer, 1x10^5^ Th17 cells were either transferred alone or in combination with 2.5x10^3^ Foxp3^+^ Treg cells or 5x10^5^ Tr1 cells into *Rag1^-/-^* mice. This transfer was performed 24 h prior to the induction of crescentic glomerulonephritis.

### Morphological examination

Light microscopy was performed on paraffin-embedded, 1 µm thick, 4 % PFA fixed kidney cross-sections stained with PAS-staining. The formation of crescents was assessed in 30 glomeruli per mouse in a blinded manner.

### Isolation of lymphocytes from different organs

Spleens and lymph nodes were smashed through a 100 µm cell strainer and washed with PBS/1 % FBS followed by centrifugation. Spleens were further processed for erythrocyte lysis using Ammonium-Chloride-Potassium- (ACK) buffer.

Kidneys were crushed and incubated at 37°C for 45 minutes in RPMI medium, containing 10 % FBS, 200 µg/ml DNase I and 2 mg/ml Collagenase D. After incubation, the tissue was mechanically reduced to a pulp, and cells were pelleted. A 37 % Percoll gradient centrifugation was performed. The remaining erythrocytes were lysed via ACK buffer.

For the isolation of intraepithelial lymphocytes (IELs), 0.5 cm pieces of the small intestines were incubated in 10 ml RPMI medium containing 10 % FBS and 1.5 mg DTT for 20 minutes at 37°C. For the isolation of lamina propria lymphocytes (LPLs), digested tissue was cut to a pulp and transferred in 6 ml Collagenase solution (RPMI medium containing 10 % FBS, collagenase (100 U/ml), and DNAse I (1000 U/ml)) for 45 minutes at 37°C. LPLs were pelleted and pooled with IELs. For further cell separation, a 40 %/ 67 % Percoll gradient was performed (GE Healthcare). The leukocyte-enriched interphase was isolated, washed, and processed for further staining steps.

### Flow cytometry

In the case of reporter mice, cells were analyzed after surface staining. To enable intracellular staining, in the case of non-reporter mice, cells were stimulated for 3 hours at 37°C with PMA (50ng/ml; Merck Darmstadt) and Ionomycin (1 mM; Sigma Aldrich).

To discriminate dead from living cells, isolated cells were pelleted and stained with a fluorochrome-labeled violet DNA dye. Pacific OrangeTM Succinimidyl Ester (Life technologies, Lot 2179293) was diluted (1:1000) in PBS and incubated for 30 minutes at 4°C. Cells were washed and pelleted.

Surface staining panel for flow cytometric analysis: CD11b (PE-Cy7, BioLegend, clone M1/70, Lot B249268, Dilution 1:400); CD11c (PE-Cy7, BioLegend, clone N418, Lot B222652, Dilution 1:400); CD195 (CCR5) (PE/Cy7, BioLegend, clone HM-CCR5, Lot B224462, Dilution 1:400); CD25 (BV650, BioLegend, clone PC61, Lot B288551, Dilution 1:100); CD3 (BUV379, BD, clone 17A2, Lot 9080908, Dilution 1:200); CD4 (Pac Blue, BioLegend, clone RM4-5, Lot B336509, Dilution 1:600; BV650, BioLegend, clone RM4-5, Lot B297638, Dilution 1:400); CD45 (BV785, BioLegend, clone 30-F11, Lot B336128, Dilution 1:800); CD45.1 (APC, BioLegend, clone A20, Lot B209251, Dilution 1:400); CD45.2 (PE Cy7, BioLegend, clone 104, Lot B307583, Dilution 1:400); CD8α (PE-Cy7, BioLegend, clone 53-6,7, Lot B295389, Dilution 1:400; NK1.1 (PE-Cy7, BioLegend, clone PK136, Lot B284829, Dilution 1:400); CD279 PD1 (BV 605, BioLegend, clone 29F.1A12, Lot B227579, Dilution 1:400); TCR-γδ (PE-Cy7, BioLegend, clone GL3, Lot B222125, Dilution 1:400); TIGIT (PerCPCy5.5, eBioscience, clone GIGD7, Lot 1988570, Dilution 1:400); CD366 (TIM-3) (BV 421, BioLegend, clone RMT3-23, Lot B259713, Dilution 1:400). Surface staining was performed for 20 minutes at 4°C.

In the case of CD49b (PE, BioLegend, clone HMa2, Lot B230820, Dilution 1:100) and CD223 (LAG3) (APC, BioLegend, clone C9B7W, Lot B243438, Dilution 1:100), the staining was performed for 30 minutes at 37°C.

In order to stain for intracellular markers, cells were fixed for 20 minutes with 4 % formaldehyde solution, followed by permeabilization of the cell membranes by adding 0.1 % Nonidet P40 (NP40)- solution (Sigma Aldrich) for 4 minutes. Lastly, a staining cocktail with fluorochrome-labeled antibodies for intracellular staining was added to the cells.

Intracellular staining panel for flow cytometric analysis: Foxp3 (APC, eBioscience, clone FJK-16s, Lot 2297415, Dilution 1:80; PE, eBioscience, clone NRRF-30, Lot 1927456, Dilution 1:80); IFN-γ (BV785, BioLegend, clone XMG1.2, Lot B343101, Dilution 1:100; APC, Biolegend, clone XMG1.2, Lot B288855, Dilution 1:100); IL-10 (PE-Dazzle, BioLegend, clone JES5-16E3, Lot B265048, Dilution 1:100); IL-17A (BV 421, Biolegend, clone TC11-18H10.1, Lot B318238, Dilution 1:100). Intracellular staining was performed for one hour at room temperature.

Fluorochrome detection was performed on an LSR II flow cytometer using FACS Diva software. For analysis, data were exported from the FACS Diva to FlowJo vX software for MAC or Windows.

### *In vitro* suppression assay

Using MACS beads, responder cells (CD4^+^ CD25^-^) were isolated from the spleen and lymph nodes and labeled with 5 µM CellTrace violet dye. Per well 1.5x10^4^ responder cells together with 7.5x10^4^ APCs were plated in a 96-well round bottom plate. Next, 1x10^4^ Treg cells/well were added to the responder-APC-mix. Additional soluble anti-CD3 specific antibody (1.5 mg/ml) led to further cell stimulation. After 96 hours at 37°C in a 5 % CO_2_ incubator, the proliferation of responder cells was determined via flow cytometry by detecting the intensity of the violet dye per cell.

### RNA isolation from sorted kidney cells to perform 10X single-cell sequencing

After sorting of 6x10^4^ Foxp3^neg^ IL-10 producing CD4^+^ T cells from nephritic kidneys, the sample was processed with Chromium TM Single Cell 3’ v2 kit according to the corresponding protocol of 10X Genomics^TM^. The libraries were sequenced on an Illumina NovaSeq 6000 system (S4 flow cell) with 150 base pairs and paired-end configurations.

### Data analysis of single-cell sequencing

Mice single-cell RNA sequencing data were processed using CellRanger version 4.0.0. The further processing and downstream analysis of the single-cell data were done using the R software version 4.1.1 (2021-08-10). The global seed was set to 0. Unless mentioned otherwise, methods were run with default parameters. The R-Package Seurat (version 4.1.0) was used for pre-processing, dimensional reduction, and cluster identification.

As part of the quality check, cells with number of genes between >450 <4000 (control) or >700 <4000 (anti-CD3) were kept, and with percentage of mitochondrial genes <35 %. Values were determined after visually inspecting the distribution. The two mouse datasets (control and anti-CD3) were integrated, scaled (with regression of library size and percentage of mitochondrial genes) and principal components (PC) were computed according to Seurat’s pipeline. After Louvain clustering (PC 1 to 30, resolution 0.8) and UMAP visualization (PC 1 to 30), clusters owing less than 150 cells as well as one cluster expressing primarily ribosomal genes and another cluster expressing cell cycle genes were removed. Finally, 10,802 cells (control 3,142, anti-CD3 7,660) were further analyzed. Next, re-clustering (resolution 0.4 and PC 1 to 25) was performed and a new UMAP was computed.

The preprocessed and clustered human data was received from (25). The dataset was subset using only CD4 positive clusters, annotated with Th1, Th17, CD4_Tcm, and Treg. Subsequently, the patients were re-integrated using methods FindIntegrationAnchords (k.filter=50), IntegrateData (k.filter = 50), and cells were re-clustered (resolution 0.5, PC 1 to 30).

Tr1 or CIR scores were calculated using Seurat’s method AddModulScore and the following mouse (or human equivalent) genes: *Il10*, *Lag3*, *Havcr2*, *Pdcd1*, *Ctla4*, *Itga2*, and *Tigit*. Significance between datasets was determined using the Wilcoxon rank sum test.

### Statistical analysis

Statistical evaluations were performed using GraphPad Prism software. Unpaired, nonparametric Wilcoxon-Mann-Whitney test, One-way ANOVA using Tukey’s multiple comparisons or Kruskal-Wallis using Dunn’s multiple comparisons test were used for the evaluation of statistical significance. A p-value < 0.05 was used to define significance.

## Results

### Analysis of IL-10-producing T-cell populations in crescentic glomerulonephritis

As a first step, we wanted to analyze the infiltration of IL-10-producing CD4^+^ T cells during the time course of crescentic glomerulonephritis (cGN). To that end, we used Fate+ reporter mice, which allow us to analyze Foxp3, IL-10, and IL-17A expression simultaneously. Furthermore, as IL-17A^+^ cells are permanently marked with YFP, we can also identify IL-10-producing cells that have previously expressed IL-17A (21). Experimental cGN was induced in Fate+ mice and we analyzed the kidneys throughout the course of the disease (Fig. 1a-c). As a control, we analyzed the frequency of these cells under steady-state conditions (day 0). IL-10 expression was detected in Foxp3^+^ (Fig. 1d+e) as well as in Foxp3^neg^ CD4^+^ T cells (Fig. 1d+f). Both IL-10-producing T-cell subsets were below 1% of the total CD4^+^ T-cell population at steady-state, but frequencies increased with the onset of the disease at day 3 reaching the peak at day 7-10 (Fig. 1e+f). However, within CD4^+^ T cells, the frequencies of Foxp3^neg^ IL-10-producing CD4^+^ T cells was about ten times higher compared to Foxp3^+^ IL-10-producing CD4^+^ T cells (Figure 1e+f). We furthermore found that a small fraction (1–5 %) of Foxp3^neg^ IL-10-producing CD4^+^ T cells were YFP^+^, indicating that they were Th17 cells or had expressed IL-17A before (Fig. 1g). Taken together, Foxp3^neg^ IL-10-producing CD4^+^ T cells emerge during the course of crescentic glomerulonephritis. Likewise, Foxp3^+^ IL-10-producing CD4^+^ T cells increase but overall at about 10 times lower levels compared to Foxp3^neg^ IL-10- producing CD4^+^ T cells. Finally, about 5% of Foxp3^neg^ IL-10-producing CD4^+^ T cells at some point produced IL-17A.

**Figure 1.**
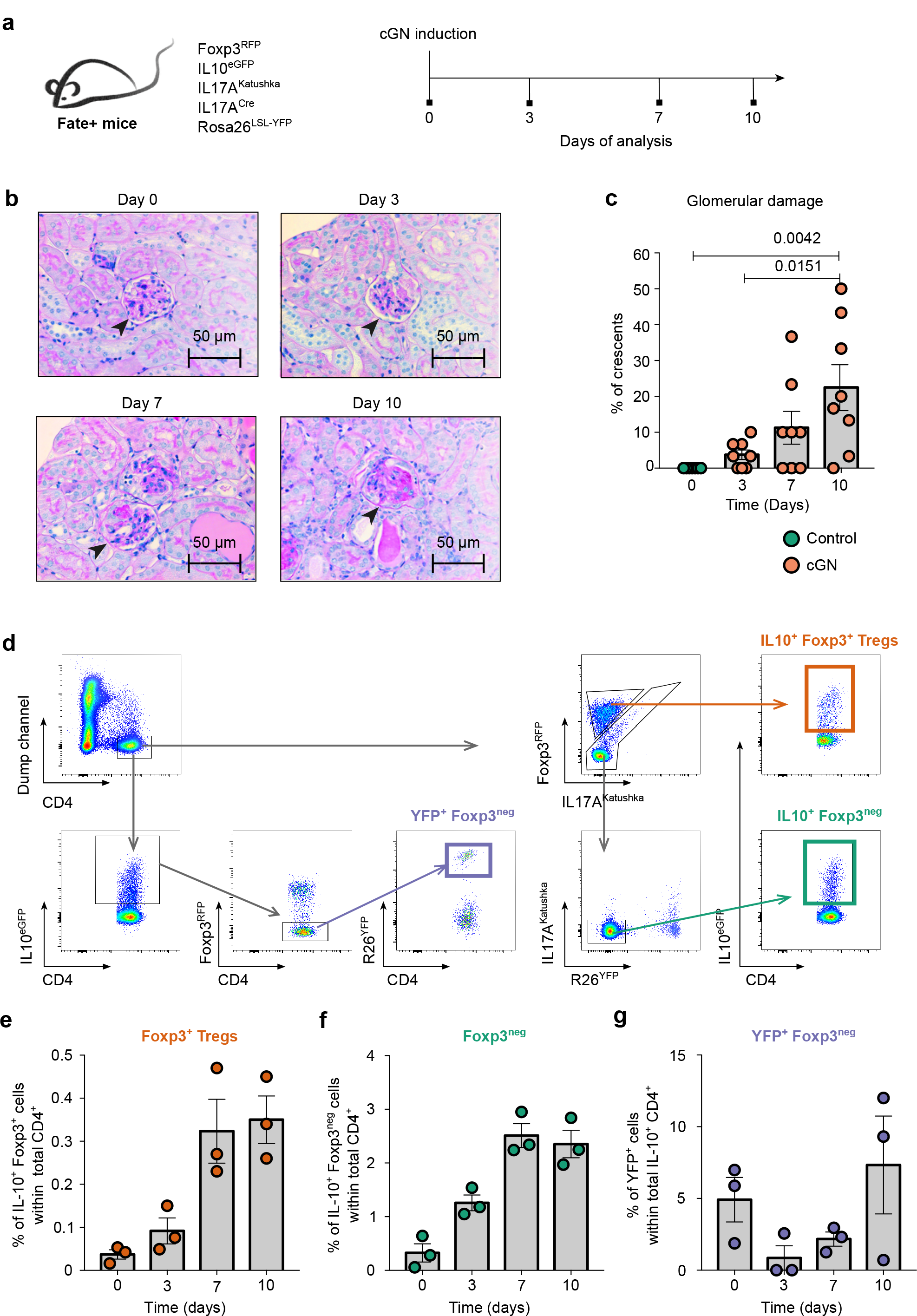
Emergence of IL-10-producing T cells in glomerulonephritis **a)** Experimental cGN was induced in Foxp3^mRFP^ IL10^eGFP^ IL17A^Katushka^ IL17A^Cre^ Rosa26^YFP^ (Fate+) mice. Animals were sacrificed under steady-state conditions and on days 3, 7, and 10 after disease induction. Cells were isolated from the kidneys. **b)** Representative PAS staining of Paraffin-embedded kidney cross sections. **c)** Percentage of crescent formation. **d)** Representative dot plots and gating strategy. **e-f)** Bar graphs depicting the percentage of **(e)** IL-10^+^ Foxp3^+^ Tregs and **(f)** IL-10^+^ Foxp3^neg^ cells within total CD4^+^ T cells. **g)** Bar graph depicting the percentage of YFP^+^ Foxp3^neg^ cells within IL-10^+^ CD4^+^ T cells. Data in (c) are representative of two independent experiments. Day 0 n=7; day 3 n=8; day 7 n=8; day 10 n=8. For statistical analysis, a One-way ANOVA with Tukey’s multiple comparisons test was used. Statistical significance was set at p<0.05. Data in (e-g) are representative of three independent experiments. Day 0 n=3; day 3 n=3; day 7 n=3; day 10 n=3. Lines indicate mean ± S.E.M.

### CD3-specific antibody treatment promotes IL-10-producing T cells in glomerulonephritis

Next, we aimed to study whether we can further expand Foxp3^neg^ IL-10-producing CD4^+^ T cells in the kidney. To this end, we used anti-CD3-specific antibody treatment, which we have used before to suppress cGN (1), and which is known to induce IL-10 production in CD4^+^ T cells in the intestine (11, 17, 21, 24). Again, we induced experimental cGN in Fate+ mice which were treated with an antibody against CD3 on days 8 and 10 post-disease induction (Fig. 2a). Kidney CD4^+^ T cells were separated into IL-10-producing Foxp3^+^ (Fig. 2b+c) and Foxp3^neg^ cells. Foxp3^neg^ IL-10-producing CD4^+^ T cells were further separated into Foxp3^neg^ YFP^neg^ cells (IL- 10-producing cells, which did not emerge from Th17 cells) (Fig. 2b+d), Foxp3^neg^ YFP^+^ IL-17A^neg^ (exTh17 cells; IL-10-producing cells, which have produced IL-17A before) (Fig. 2b+e) and Foxp3^neg^ YFP^+^ IL-17A^+^ (IL-10-producing Th17 cells) (Fig. 2b+f). In line with previous data studying the intestine (Fig. S1), anti-CD3-antibody treatment strongly increased the frequency of IL-10 in all analyzed subsets in the kidney (Fig. 2c-f). Furthermore, we confirmed that some exTh17 cells expressed IL-10. Also, here the production of IL-10 was further promoted by anti-CD3-antibody treatment (Fig. 2e).

**Figure 2.**
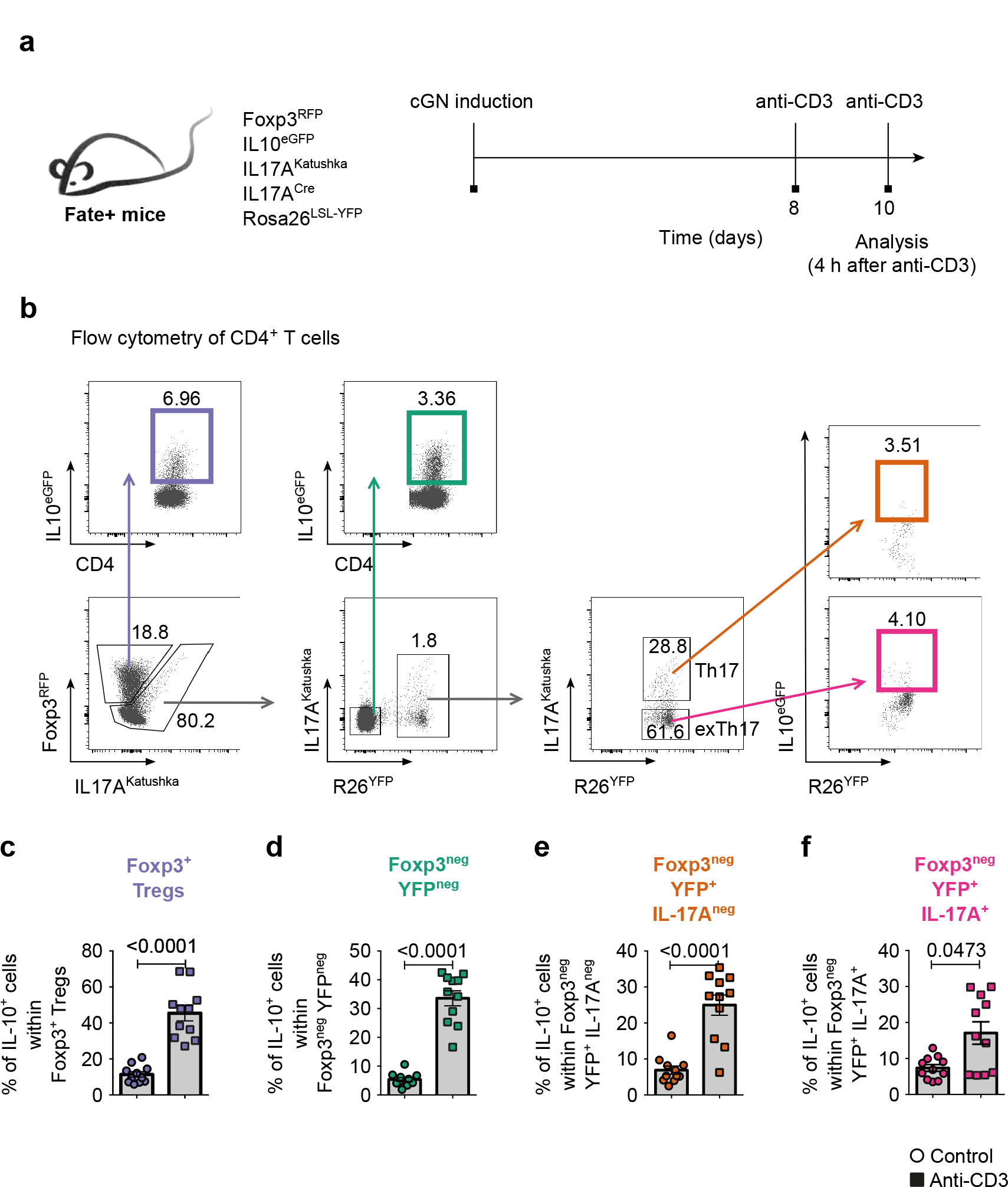
Expansion of IL-10-producing T cells in glomerulonephritis upon CD3-specific antibody treatment **a)** Experimental cGN was induced in Foxp3^mRFP^ IL10^eGFP^ IL17A^Katushka^ IL17A^Cre^ Rosa26^YFP^ (Fate+) mice separated into two groups. One group received additional 15 µg of anti-CD3 specific antibody on days 8 and 10 post disease induction. Control group received PBS. Mice were sacrificed 4 hours after the last antibody or PBS injection. Cells were isolated from the kidneys. **b)** Representative dot plots and gating strategy. **c-f)** Bar graphs depicting the percentages of IL-10^+^ cells within **(c)** Foxp3^+^ Tregs, **(d)** Foxp3^neg^ YFP^neg^, **(e)** Foxp3^neg^ YFP^+^ IL17A^neg^, and **(f)** Foxp3^neg^ YFP^+^ IL17A^+^ cells. Data in (c-f) are cumulative of four independent experiments. Control n=11; anti-CD3 n=11. For statistical analysis a Wilcoxon-Mann-Whitney test was used. Statistical significance was set at p<0.05. Lines indicate mean ± S.E.M.

### Single-cell sequencing reveals that a fraction of Foxp3^neg^ IL-10-producing CD4^+^ T cells in cGN expresses the core transcriptional program of Tr1 cells

Next we aimed to further characterize Foxp3^neg^ IL-10-producing CD4^+^ T cells in the kidney in an unsupervised approach. To this end, we used single-cell sequencing. We induced experimental cGN in Foxp3^RFP^ IL10^eGFP^ reporter mice and treated half of them with an antibody against CD3. Control group received PBS instead. Foxp3^neg^ IL-10-producing CD4^+^ T cells were FACS sorted and then subjected to single-cell sequencing using the 10X genomics platform (Fig. 3a and S2a). A total of 10,802 cells (control: 3,142 cells, anti-CD3: 7,660 cells) were analyzed. This analysis revealed a heterogeneous cell population that separated into seven distinct clusters (C1-C7) (Fig. 3b). While most of the clusters showed a gene signature towards effector-like cells, clusters 3 and 4 were enriched with CIR (Fig. 3b+c). The expression of specific genes within the clusters can be viewed in Fig. S2b. Moreover, we specifically focused on the identification of Tr1 cells, and screened the genes to identify them. This analysis also revealed clusters 3 and 4 to express the highest Tr1 score (combination of *Il10*, *Lag3*, *Havcr2*, *Pdcd1*, *Ctla4*, *Itga2*, and *Tigit*) (Fig. 3d). Additionally, the treatment with anti-CD3 specific antibody increased the frequency of co-inhibitory receptors (CIR)-expressing Foxp3^neg^ IL-10-producing CD4^+^ T cells. On this basis, we next looked at the clusters from the control and anti-CD3 treated groups separately and checked for the relative proportion of each cluster (Fig. 3c). Here, the additional treatment with an antibody against CD3 led to a relative shrink of clusters 1, 2, 6, and 7 but a relative increase of clusters 3, 4, and 5 (Fig. 3c). Moreover, also the Tr1 score was increased in cells derived from mice treated with anti-CD3 antibody (Fig. 3e). As expected, when differentially expressed genes were plotted comparing clusters 3 and 4 versus the rest, many genes such as *Lag3*, *Ctla4*, *Pdcd1*, and *Havcr2*, responsible for the *bona fide* Tr1 signature emerged (Fig. 3f).

**Figure 3.**
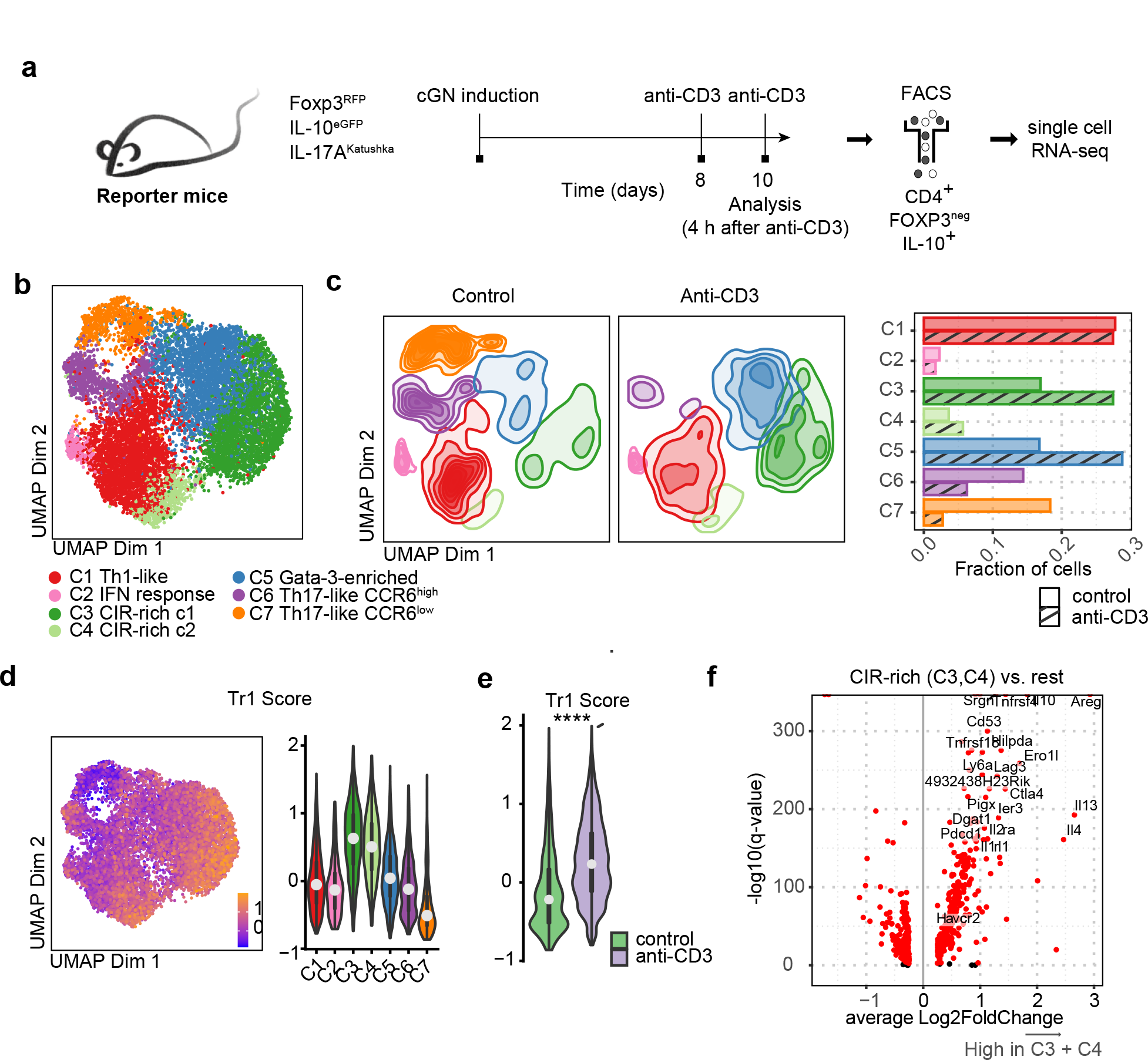
A subset of IL-10-producing T cells in the kidney expresses the core transcriptional program of Tr1 cells **a)** Experimental cGN was induced in Foxp3^mRFP^ IL10^eGFP^ IL17A^Katushka^ mice. On days 8 and 10 after disease induction mice were injected either with PBS or anti-CD3 antibody. Cells were isolated 4 h after the last injection from the kidneys of nephritic mice. **b)** UMAP and cluster identification within IL-10^+^ CD4^+^ T cells. **c)** UMAP and relative frequency of clusters 1-7 from control group and anti-CD3-treated mice. **d)** Heat-UMAP and Violin plot of Tr1 signature-score in clusters 1-7 in control group and anti-CD3-treated mice. **e)** Tr1 score comparison in control and anti-CD3-treated groups. Tr1 score is based on the expression of: *Il10, Lag3, Havcr2, Pdcd1, Ctla4, Itga2, Tigit*. **f)** Volcano plot of differentially expressed genes comparing clusters 3 and 4 versus the rest. For statistical analysis a Wilcoxon rank sum test was used. **** p<0.0001.

Taken together, we identified Foxp3^neg^ IL-10-producing CD4^+^ T cells in murine nephritic kidneys, which express the core transcriptional program of Tr1 cells (e.g. cluster 3 and 4). This fraction was further expanded upon anti-CD3 specific antibody treatment.

### Tr1 cells can be identified by the surface expression of co-inhibitory receptors in cGN

Having identified CD4^+^ T cells expressing the characteristic genes of Tr1 cells in the kidneys of nephritic mice, we then focused on identifying these cells via flow cytometry. Therefore, we specifically analyzed the expression of CIR which in the single-cell sequencing analysis were shown to be mainly expressed by Tr1 cells, essential as shown by us before in the intestine (17). For this, we induced cGN in Foxp3^RFP^ IL10^eGFP^ reporter mice. On days 8 and 10, half of the mice received an injection of an antibody against CD3. Control mice received PBS injections instead (Fig. 4a). As shown before, treatment with anti-CD3-antibody increased the percentage of Foxp3^neg^ cells that expressed IL-10 (Fig. 4b). To identify CIR-rich cells within Foxp3^neg^ IL-10-producing CD4^+^ T cells, we analyzed the co-expression of IL- 10 and CD49b, LAG-3, TIGIT, and TIM3 via flow cytometry (Fig. S3). We identified Foxp3^neg^ IL-10-producing CD4^+^ T cells that indeed additionally expressed CIR either alone or in combination with other markers (Fig. 4c-e). The most common combination at steady-state corresponded to CD49b, LAG-3, TIM3, and TIGIT. In correlation with the results from Figure 3, anti-CD3 antibody treatment also strongly induced the frequency of CIR (Fig. 4d+e). The four-marker combination was again the most expanded, followed by the combination of CD49b, LAG-3, and TIM3 and then the double combination of CD4b and LAG-3. Other combinations also increased their proportion under the anti-CD3 treatment (Fig. 4e).

**Figure 4.**
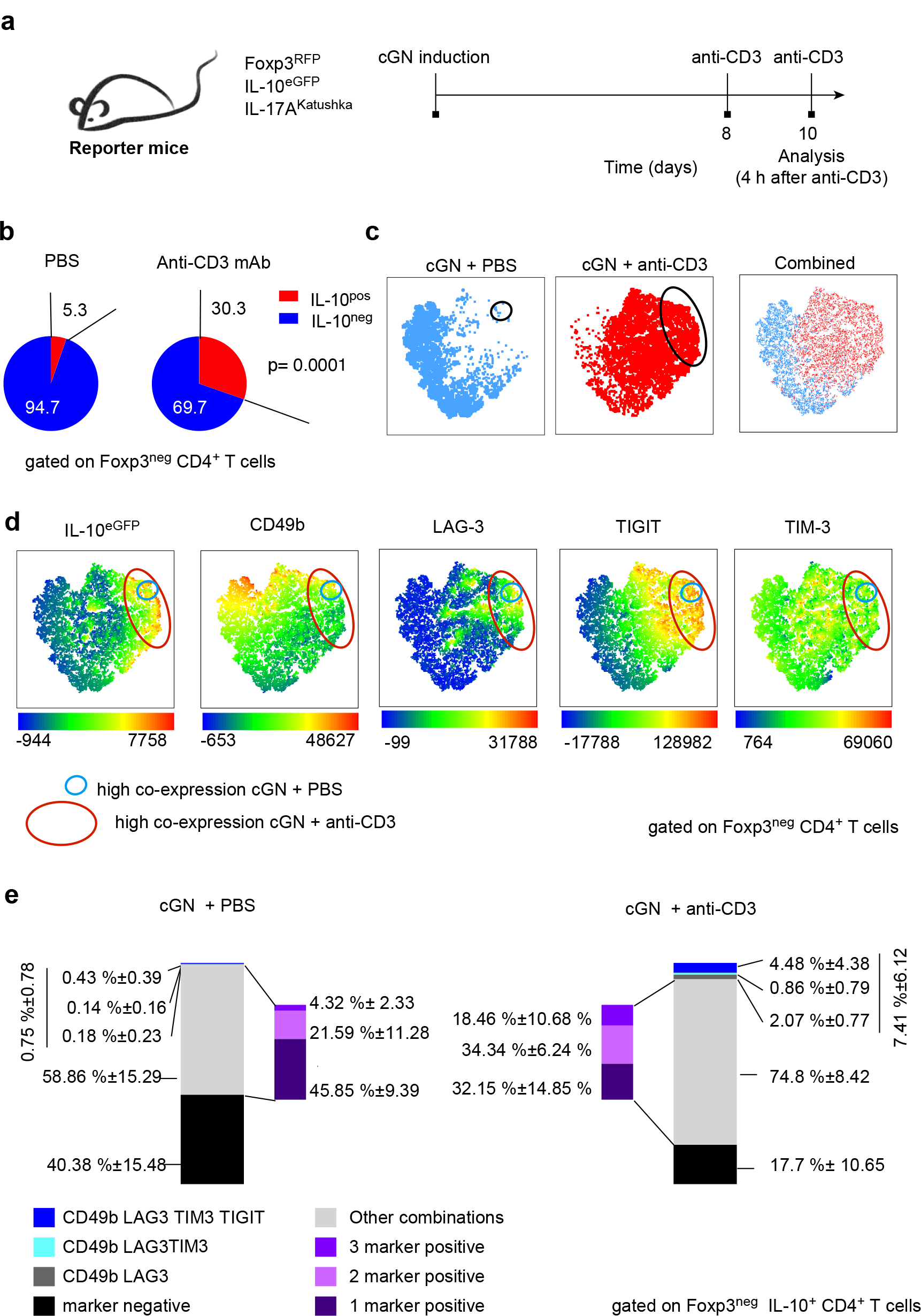
Identification of Tr1 cells within the IL-10-producing T cell subset on the basis of extra cellular markers **a)** Experimental cGN was induced in Foxp3^mRFP^ IL10^eGFP^ IL17A^Katushka^ mice. On days 8 and 10 after disease induction mice were injected either with PBS or anti-CD3 antibody. Cells were isolated 4 h after the last injection from the kidneys of nephritic mice. Analysis of IL-10 and CIR (CD49b, LAG-3, TIM-3, TIGIT) in Foxp3^neg^ CD4^+^ T cells was performed. **b)** Pie chart depicting the percentage of IL-10^+^ vs. IL-10^neg^ Foxp3^neg^ CD4^+^ T cells. **c)** tSNE graphs showing the Foxp3^neg^ CD4^+^ population in PBS vs. anti-CD3 conditions. The circles mark the position of Tr1 cells. **d)** tSNE graphs showing the expression of IL-10 or co-stimulatory surface markers gated on Foxp3^neg^ CD4^+^ T cells. The circles mark the position of Tr1 cells. **e)** Bar charts showing the distribution of stated marker combinations gated on Foxp3^neg^ IL-10- producing CD4^+^ T cells. Data in (b-e) are representative of three independent experiments. PBS n= 11; anti-CD3 n=6. For statistical analysis a Wilcoxon-Mann-Whitney test was used. Statistical significance was set at p<0.05.

Thus, anti-CD3 specific antibody treatment induced Foxp3^neg^ IL-10-producing CD4^+^ T cells. These cells represent a heterogenous population some of which express CIR, the extra-cellular markers characteristics of Tr1 cells.

### Kidney-derived IL-10-producing T cells show suppressive activity *in vitro*

Based on the above-mentioned data, we next aimed to evaluate the suppressive capacity of kidney-derived Foxp3^neg^ IL-10-producing CD4^+^ T cells. To this end, we first used an *in vitro* assay. CD4^+^ CD25-depleted cells (responder cells) were isolated from the spleens of mice at steady state and stained with a proliferation dye. During cell division, the intensity of the dye is halved in both daughter cells. To obtain IL-10-producing T cells, experimental cGN was induced in Foxp3^RFP^ IL-10^eGFP^ reporter mice and Foxp3^+^ CD4^+^ T cells, as well as Foxp3^neg^ IL-10-producing CD4^+^ T cells, were sorted out from the kidneys 10 days after disease induction and their suppressive function assessed *in vitro* (Fig. 5a). As a negative control, regulatory T cells were exchanged with the same numbers of non-suppressive CD4^+^ responder cells. The dye was measured after five days using flow cytometry (Fig. 5b). When the violet dye intensity was analyzed in the responder cells, the negative control did not show any suppression but rather enhanced proliferation of responder cells (Fig. 5b-c). The addition of kidney-derived Foxp3^neg^ IL-10-producing CD4^+^ T cells inhibited proliferation of almost 85 % of the responder cells, which was comparable to the one observed by Foxp3^+^ Tregs (Fig. 5b-c).

**Figure 5.**
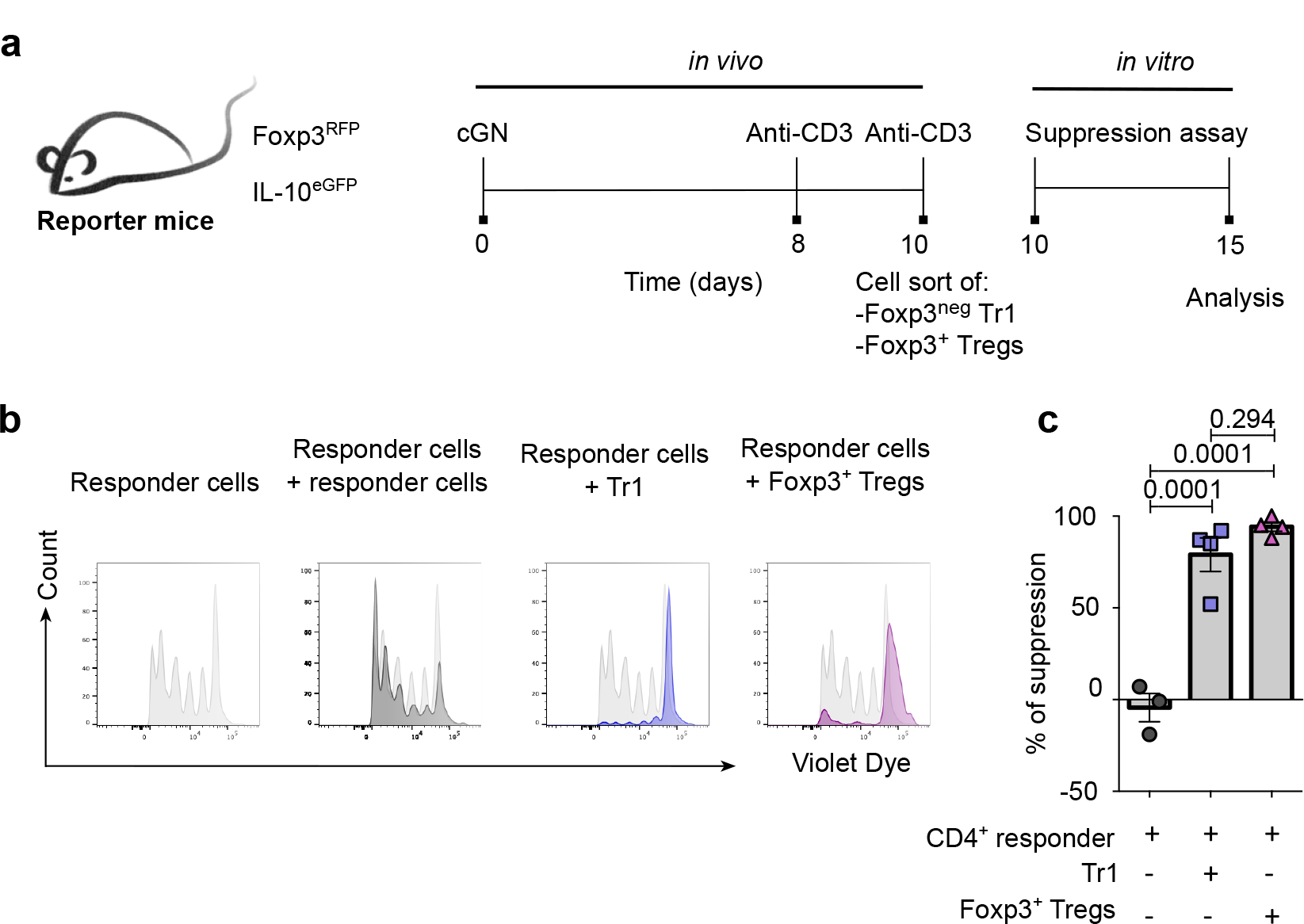
Tr1 cells from the kidney have *in vitro* suppressive activity **a)** Experimental cGN was induced in Foxp3^mRFP^ IL10^eGFP^ mice. On days 8 and 10 after disease induction mice were injected either with PBS or anti-CD3 antibody. Suppressor cells were isolated 4 h after the last injection from the kidneys of nephritic mice. 1.5 x 10^4^ CD4^+^ T cells (responder cells) were plated per well. Additional 7.5 x 10^4^ irradiated antigen presenting cells (APCs) served the purpose of cell activation. 1x10^4^ suppressor cells were added to each well. CellTrace violet dye intensity was measured via flow cytometry. **b)** Representative histograms showing the suppression of responder cells, Tr1 cells, and Foxp3^+^ Tregs. **c)** Bar graph depicting the percentage of suppression for each condition. Data in (b-c) are representative of three independent experiments. Responder + responder n=3; responder + Tr1 n=4; responder + Foxp3^+^ Tregs n=4. For statistical analysis, a One- way ANOVA with Tukey’s multiple comparisons test was used. Statistical significance was set at p<0.05. Lines indicate mean ± S.E.M.

Taken together, *ex vivo* isolated Tr1 cells from the kidney could suppress the proliferation of CD4^+^ responder T cells *in vitro* to a similar degree as classical Foxp3^+^ Tregs.

### *In vitro*-generated Tr1 cells suppress cGN

Tr1 cells are known to be efficient regulators of CD4^+^ T-cell-driven inflammation in the intestine (11, 17). Thus, having identified the presence of *bona fide* Tr1 cells within the IL-10^+^ Foxp3^neg^ population in the kidneys of nephritic mice, we next aimed to evaluate their suppressive capacity in the context of cGN. Of note, Th17 cells have been shown to be pathogenic on their own and promote glomerulonephritis when transferred into these mice (1, 5). Thus, we designed an *in vivo* suppression assay by transferring Th17 cells plus Tr1 cells into *Rag1^-/-^* mice (Fig. 6a). Since a high cell number is required for these experiments, both Th17 and Tr1 cells were differentiated *in vitro*. Foxp3^+^ Treg cells isolated from the spleens of untreated mice at steady state were used as a positive control (Fig. 6b). 24 hours after the T cell transfer, glomerulonephritis was induced in the mice, and the survival rate and disease severity were evaluated (Fig.6c). Though not significant, mice that received only Th17 cells showed a relative increase in mortality compared to mice that received additional regulatory cells (Fig. 6d). Also, the glomerular damage analyzed in kidney cross sections, revealed the highest score in mice that received Th17 cells alone (Fig. 6e-f). Both groups that received either Tr1 cells or Foxp3^+^ Tregs cells displayed not only increased survival but also less crescent formation (Fig. 6d-f).

**Figure 6.**
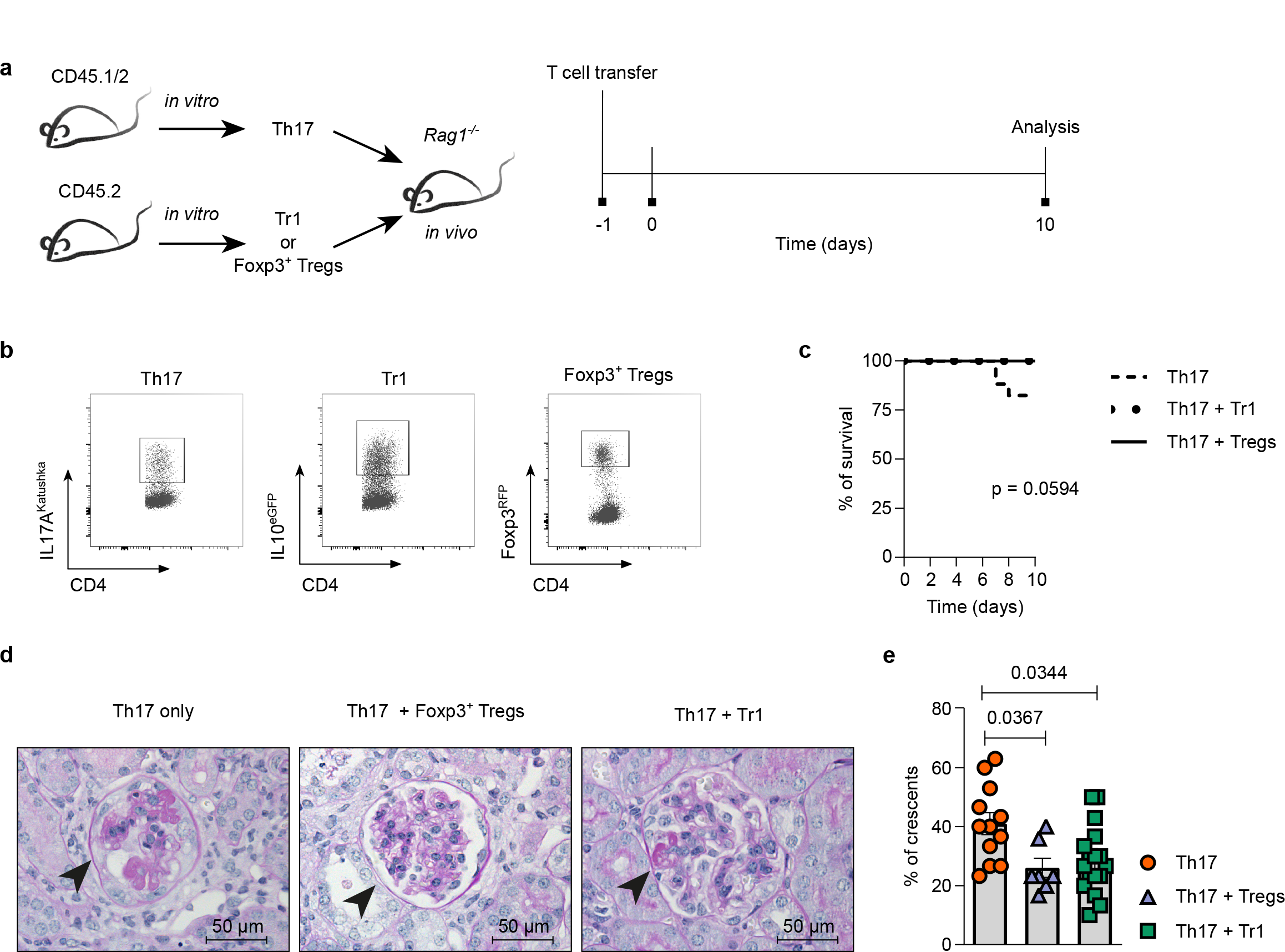
Tr1 cells suppress experimental glomerulonephritis *in vivo* **a)** Th17 and Tr1 cells were differentiated *in vitro* from CD45.1/2 or CD45.2 mice, respectively. Foxp3^+^ Tregs cells were isolated from spleens of CD45.2 mice at steady state. After differentiation, cells were sorted and Th17 cells were transferred into *Rag1^-/-^* mice either alone or in combination with Tr1 cells or Foxp3^+^ Tregs as control. Experimental cGN was induced 24 h after the cell transfer. Animals were sacrificed 10 days after disease induction. **b)** Representative dot plots and gating strategy of the sorted T cell populations. **c)** Survival curve. **d)** Representative PAS staining of Paraffin-embedded kidney cross sections. **e)** Percentage of crescent formation. Data in (c-e) are cumulative of five independent experiments. Th17 only n=12; Th17 + Foxp3^+^ Tregs n=7; Th17 + Tr1 n=17. For statistical analysis a Kruskal-Wallis with Dunn’s multiple comparison test was used. Statistical significance was set at p<0.05. Lines indicate mean ± S.E.M.

In conclusion, *in vitro* differentiated Tr1 cells were sufficient to suppress Th17 cell- mediated glomerulonephritis *in vivo*.

### Identification of Tr1 cells in ANCA-associated glomerulonephritis patients

Finally, we aimed to translate these findings into human biology. To this end, we took advantage of already available single-cell data from kidney biopsies isolated from ANCA patients (Fig. 7a) (25). UMAP analysis identified eight different clusters within human-derived CD4^+^ T cells from the kidney biopsies (Fig. 7b). We then looked at the expression of Foxp3, and CIR within the clusters. We specifically aimed for the identification of Foxp3^neg^ CD4^+^ T cells that express at least two surface markers identifying Tr1 cells (Fig. 7c). Through this approach, we identified a large number of cells that expressed at least two or more CIR at the same time and could correspond to *bona fide* Tr1 cells. Next, we identified cells that showed an increased CIR score that was previously defined by simultaneous expression of the genes for *LAG3*, *ITGA2*, *HAVCR2*, *PDCD1*, *TIGIT,* and *CCR5*. This score was high in clusters 2 and 5, and relatively increased in cluster 7 (Fig. 7d). By further analyzing the expression levels for genes encoding for *IL10*, *CTLA4*, *FOXP3,* and *IL2RA* we identified cluster 5 to be enriched in Foxp3^+^ Tregs (Fig. 7e). Thus, we were able to identify clusters 2 and 7, which showed low gene expression of *FOXP3* and *IL2RA*, as clusters corresponding to Tr1 cells. These results are in correlation with our previous results in mice, where we could observe a high heterogeneity in Foxp3^neg^ IL-10-producing T cells.

**Figure 7.**
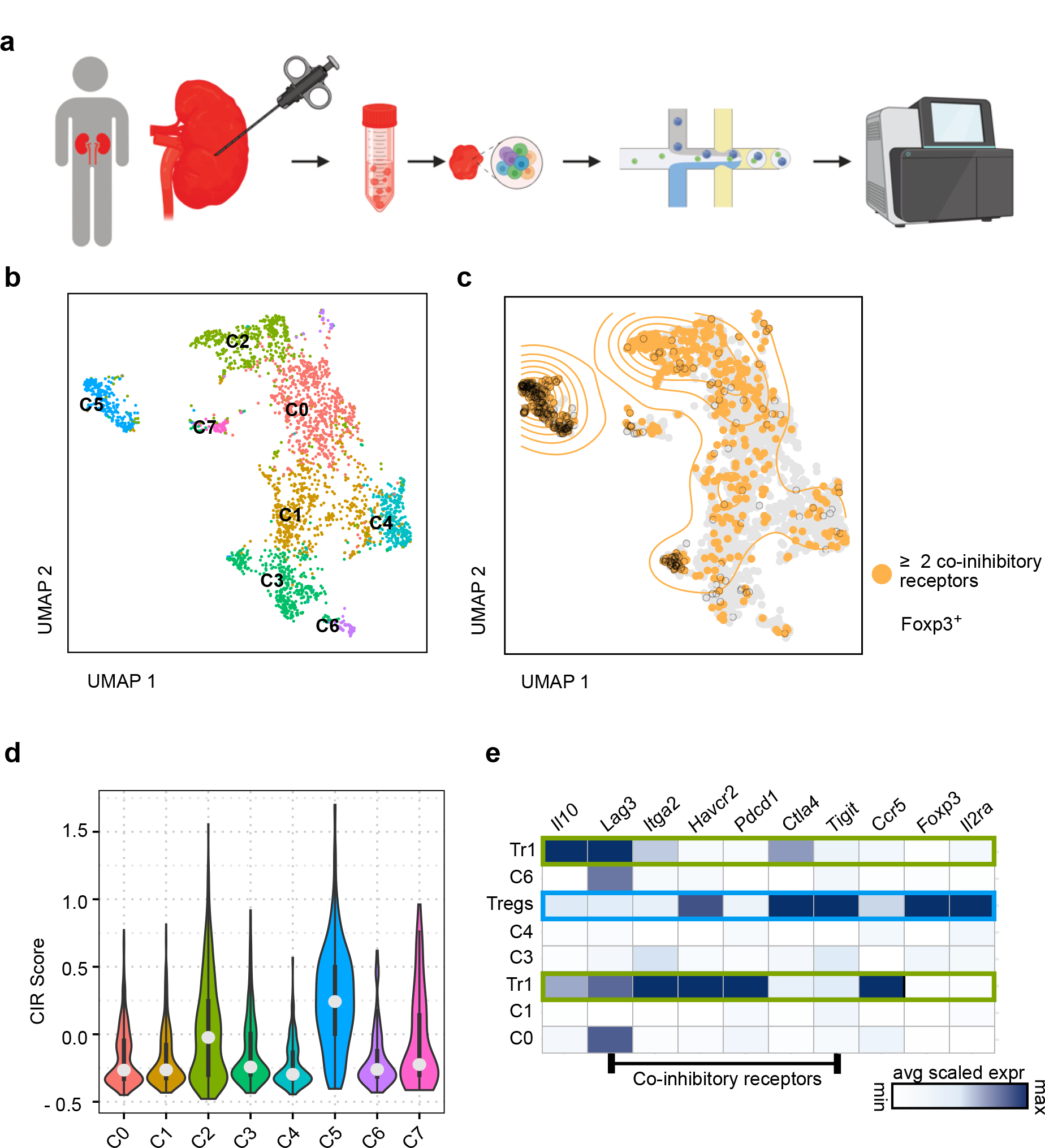
Tr1 cells in ANCA-associated glomerulonephritis patients **a)** Kidney biopsies were isolated from an ANCA patient. Biopsies were reduced to a single cell suspension. Cells were barcoded and single-cell sequencing was performed. **b)** UMAP and cluster identification within CD4^+^ T cells. **c)** UMAP of single cells expressing Foxp3 (black circles) and two or more co-inhibitory receptors (orange color) **d)** Violin-Plot of clusters 0-7 indicating the CIR-score in each cluster. **e)** Heat map of expression of indicated genes in clusters 0-7.

These data indicate that *bona fide* Tr1 cells are also present in the kidney in patients with ANCA nephritis.

## Discussion

This study aimed to address the potential of Tr1 cells to be used as T cell-based therapy for crescentic glomerulonephritis (cGN). To this end, we analyzed the emergence of Tr1 cells, identified their molecular profile, and tested their functionality in the context of glomerulonephritis. We found that Tr1 cells emerge, albeit at low numbers, during experimental glomerulonephritis, and proved their high level of suppressive activity *in vitro*. Importantly, Tr1 cell transfer was protective in a mouse cGN model. Thus, our study sets the premise to therapeutically target Tr1 cells in cGN.

Foxp3^+^ Tregs have been described as being able to ameliorate kidney injury by suppressing Th17 cell-mediated pathology via IL-10 (10). Their power to stabilize immune homeostasis was found to be common for different organs and diseases, e.g. kidneys and the intestine (11). It has been reported before that IL-10-producing Foxp3^+^ Tregs are important to suppress kidney injury in cGN (10). We here showed that Foxp3^neg^ Tregs, namely Tr1 cells, are able to produce high amounts of IL-10 and to suppress nephritis. This is in line with previous data showing that besides Foxp3^+^ Tregs, Tr1 cells can also suppress Th17 cell-driven colitis (11). Tr1 cells have already been described in other autoimmune kidney diseases (18, 19). However, in these studies, the identification of Tr1 cells was solely based on the expression of IL- 10 and the lack of Foxp3. Thus, the role of *bona fide* Tr1 cells remained to be elucidated. To answer this question, we first studied IL-10-producing CD4^+^ T cells, including Tr1 cells, in the kidney. Even under steady-state conditions, we could detect Tr1 cells in the kidney, confirming that these cells may not only be present in the intestine (26). Indeed, as the disease progressed, frequencies of IL-10 strongly increased in both Foxp3^+^ and Foxp3^neg^ CD4^+^ T cells. As mentioned before, in the disease setting the number of Foxp3^neg^ IL-10-producing CD4^+^ T cells in the kidney was about 10 times higher than the number of Foxp3^+^ IL-10-producing Tregs.

We next aimed to determine the origin of the Foxp3^neg^ IL-10-producing CD4^+^ T cells identified in the inflamed kidney. Of note, it had been shown before that Th17 cells can co-produce IL-10. Furthermore, intestinal Th17 cells were shown to be able to completely transdifferentiate into Tr1 cells (referred to as Tr1^exTh17^), a process called T cell plasticity (21). While Th17 cells in the brain of EAE mice can to convert into Th1 cells, Th17 cells in the kidney have been described to be more stable in terms of their cytokine expression and to not convert into Th1 cells to a large extent (1). To investigate whether Th17 cells found in the kidneys of nephritic mice were able to convert into Tr1 cells, we used Fate+ mice. We found that a fraction of Foxp3^neg^ IL- 10-producing CD4^+^ T cells originated from Th17 cells. Thus, we demonstrated Th17 cell plasticity towards Tr1 cells. By tracking Th17 cells, we also identified cells co- producing IL-17 and IL-10. However, the number of Tr1^exTh17^ cells was low.

To further characterize Foxp3^neg^ IL-10-producing CD4^+^ T cells in an unsupervised way, we performed single-cell RNA sequencing of cells isolated from the kidney of mice with glomerulonephritis and nephritic mice that were additionally treated with an anti-CD3 antibody. This approach confirmed that a fraction of Foxp3^neg^ IL-10- producing CD4^+^ T cells from the kidney showed the transcriptional signature of a Tr1 cell in the inflamed kidney. For both samples, the clustering of the total cells displayed a heterogeneous cell population. While some clusters revealed gene expression levels associated with effector cells, others displayed a regulatory-like gene expression pattern. This is in line with previous data showing that in the small intestine and the spleen, there was a strong heterogeneity of IL-10-producing CD4^+^ T cells (17). Recently, it was shown that IL-10 arising from CD4^+^ T cells can have a pathogenic role in EAE by promoting the survival of effector T cells (27). Though one could speculate a similar role in the kidney, especially considering the presence of pro-inflammatory-like IL-10^+^ CD4^+^ T cell clusters, it is unknown whether these cells are pathogenic in nephritis. Furthermore, in both EAE and experimental cGN a complete lack of IL-10 results in increased disease severity (10, 27). Thus, further studies will be essential to decipher the role of Tr1 cells, which suppress nephritis, from other cellular sources of IL-10, which may promote disease.

To build upon these descriptive data, we aimed to gain confirmation regarding whether kidney-derived Foxp3^neg^ IL-10-producing CD4^+^ T cells would fulfill the ultimate criterion defining Tr1 cells, i.e. regulatory activity. We performed *in vitro-* suppression assays with Foxp3^neg^ IL-10-producing CD4^+^ T cells from the kidneys of nephritic mice showing that these cells indeed have a regulatory function *in vitro*. Thus, one could envision using these cells as means of T cell-based therapy. However, to form the basis for this, one needs to further show their regulatory function *in vivo*. Indeed, during ongoing kidney inflammation, Foxp3^+^ Tregs are present and increase in numbers during disease progression (9). These cells are able to suppress CD4^+^ T cell-driven inflammation in the kidneys (9, 10, 28). Thus, we wanted to go further and determine the suppressive capacity of Tr1 cells, in an *in vivo* suppression assay combined with glomerulonephritis. To avoid confounding factors and to be able to study the interaction between pathogenic and regulatory CD4^+^ T cells only, we performed a transfer experiment of Th17 cells alone or together with Tr1 cells and Foxp3^+^ Tregs respectively into *Rag1^-/-^* prior to induction of cGN. To this end, regulatory T cells had to be first generated *in vitro* as it was not possible to obtain a sufficient number of Tr1 cells from the kidneys. Moreover, since the relative contribution of each IL-10^+^ cluster was not defined, we performed these experiments by transferring the whole IL-10^+^ Foxp3^neg^ population. Strikingly, our data indicate that in these conditions Tr1 cells are capable of suppressing Th17 cell- mediated glomerulonephritis. Thus, our data form the basis to establish a Tr1-based T-cell therapy for patients with glomerulonephritis, similar to what is already done in IBD patients (29, 30).

Finally, to further support this, we aimed to prove the existence of *bona fide* Tr1 cells in human kidneys. For this, we took advantage of available single-cell transcriptome data from ANCA patients to establish an identification method for human Tr1 cells (25). We focused on CD4^+^ T cells that expressed CIR and other surface markers. We shed light on the population of IL-10-producing CD4^+^ T cells and finally, showed the gene expression pattern of Tr1 cells. The results shown here prove that Tr1 cells are present in the human kidney. However, and in agreement with the mouse data, there is a high degree of heterogeneity in IL-10^+^ CD4^+^ T cells. The contribution of each cluster to the regulatory function or whether they represent a source of pathogenic IL-10 remains to be studied.

In conclusion, we identified Tr1 cells in mouse and human diseased kidneys. These cells have strong suppressive activity in a mouse model of cGN. Thus, our data form the basis not only to identify Tr1 cells in cGN-patients but also to potentially develop personalized T-cell therapies for patients with crescentic glomerulonephritis.

## Acknowledgements

We thank the FACS Sorting Core Facility and the Single Cell Core Facility of the Universitätsklinikum Hamburg-Eppendorf for their support. We thank Cathleen Haueis, Sandra Wende and Anett Peters for their excellent technical assistance. This work was supported by a grant from the *Deutsche Forschungsgemeinschaft* (DFG) (SFB1192 project A5 to C.F.K. and S.H.) and a grant of the *Forschungsförderungsfonds der Medizinischen Fakultät* (FFM) to S.S.-W.

## Author information

These authors contributed equally:

Shiwa Soukou-Wargalla, Christoph Kilian Christian F. Krebs, Samuel Huber

## Authors and Affiliations

I. Department of Medicine, University Medical Center Hamburg-Eppendorf, 20246, Hamburg, Germany

Shiwa Soukou-Wargalla, Christoph Kilian, Lis Velasquez, Andres Machicote, Friederike Stumme, Tanja Bedke, Jan Kempski, Morsal Sabihi, Penelope Pelczar, Beibei Liu, Can Ergen, Babett Steglich, Samuel Huber

III. Department of Medicine, University Medical Center Hamburg-Eppendorf, Hamburg, Germany

Christian F. Krebs, Hans-Joachim Paust, Ning Song, Ulf Panzer, Tobias B. Huber, Alina Borchers

Hamburg Center for Translational Immunology, University Medical Center Hamburg-Eppendorf, Hamburg, Germany

Tobias B. Huber, Ulf Panzer, Nicola Gagliani, Christian F. Krebs, Samuel Huber

Department for General, Visceral and Thoracic Surgery, University Medical Center Hamburg-Eppendorf, Hamburg, Germany

Anastasios Giannou, Nicola Gagliani, Franziska Muscate

OneChain Immunotherapeutics, University of Barcelona, Barcelona, Spain

Laura Garcia Perez

## Ethics declarations

Competing interests: The authors declare no competing interests.

## Suppl. Figures

**Figure S1.**
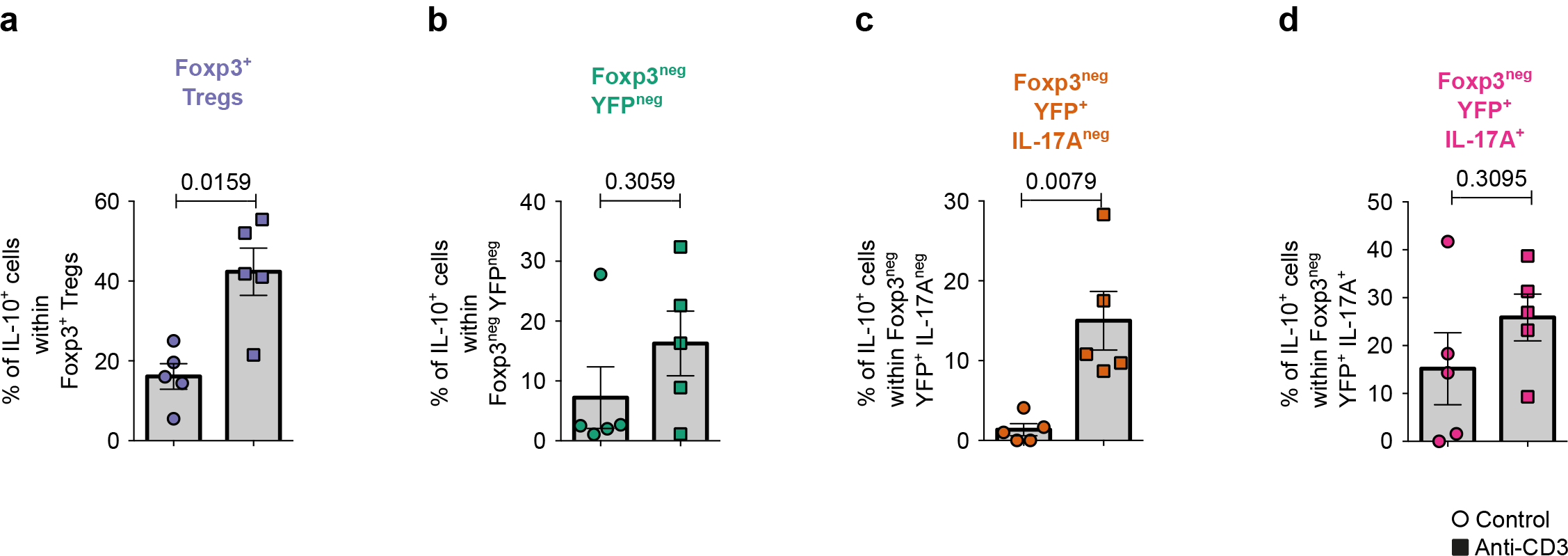
anti-CD3-specific antibody treatment induces IL-10-producing T cells in the gut of nephritic mice. Experimental cGN was induced in Foxp3^mRFP^ IL10^eGFP^ IL17A^Katushka^ IL17A^Cre^ Rosa26^YFP^ (Fate+) mice separated into two groups. One group received additional 15 µg of anti-CD3 specific antibody on days 8 and 10 post disease induction. Control group received PBS. Mice were sacrificed 4 hours after the last antibody or PBS injection. Cells were isolated from the small intestine. **a-d)** Bar graphs depicting the percentages of IL-10^+^ cells within **(a)** Foxp3^+^ Tregs, **(b)** Foxp3^neg^ YFP^neg^, **(c)** Foxp3^neg^ YFP^+^ IL17A^neg^, and **(d)** Foxp3^neg^ YFP^+^ IL17A^+^ cells. Data are cumulative of three independent experiments. Control n=5; anti-CD3 n=5. For statistical analysis a Wilcoxon-Mann-Whitney test was used. Statistical significance was set at p<0.05. Lines indicate mean ± S.E.M.

**Figure S2.**
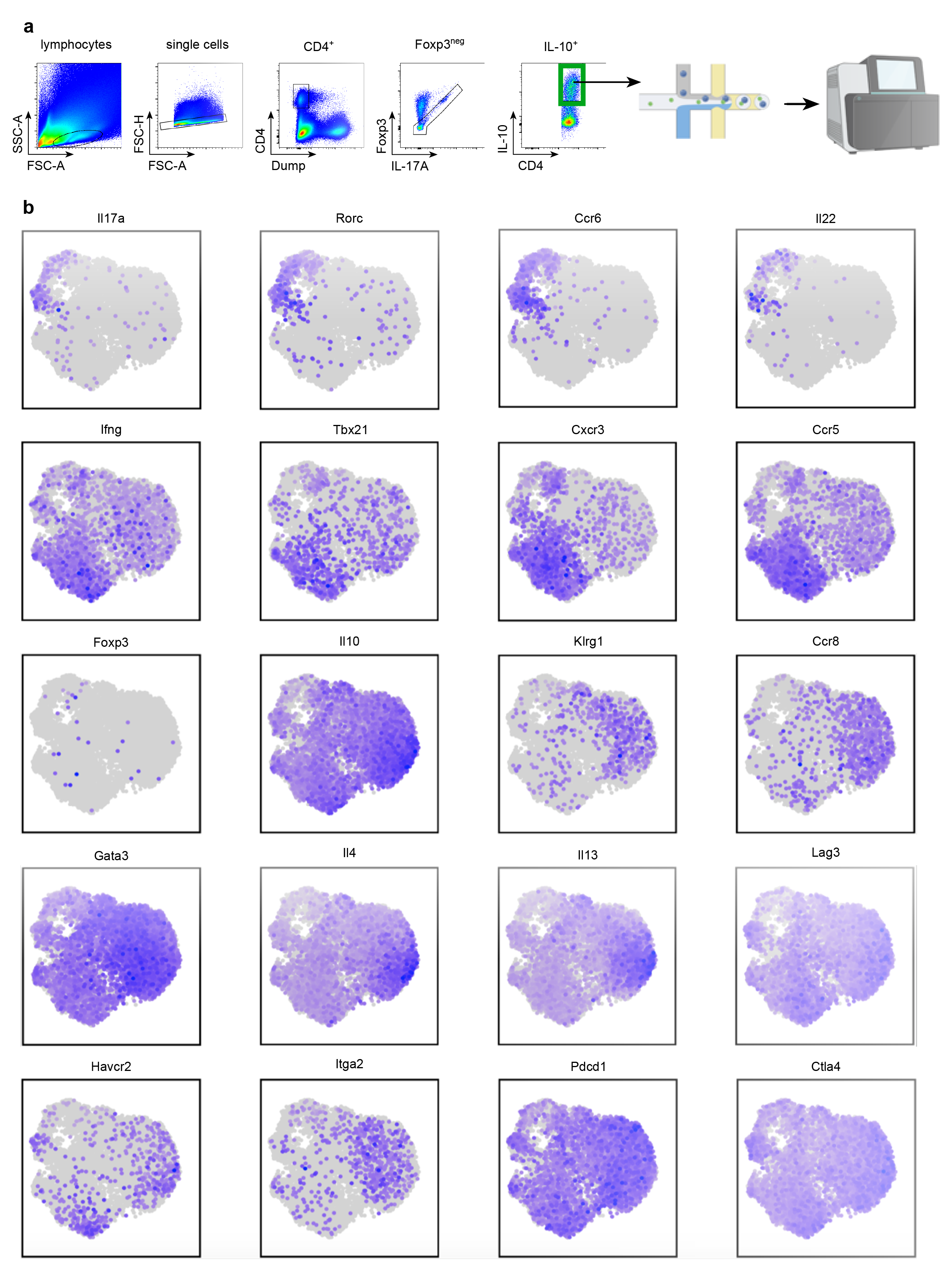
Gene expression of Tr1-associated and pro-inflammatory genes in IL-10-producing T cell clusters in the kidney **a)** Gating strategy for sorting of IL-10^+^ cells in Figure 3. **b)** UMAP depicting the gene expression pattern of *Il17a*, *Rorc*, *Ccr6*, *Il22*, *Ifng*, *Tbx21*, *Cxcr3*, *Ccr5*, *Foxp3*, *Il10*, *Klrg1*, *Ccr8*, *Gata3*, *Il4*, *Il13*, *Lag3*, *Havcr2*, *Itga2*, *Pdcd1*, and *Ctla4*.

**Figure S3.**
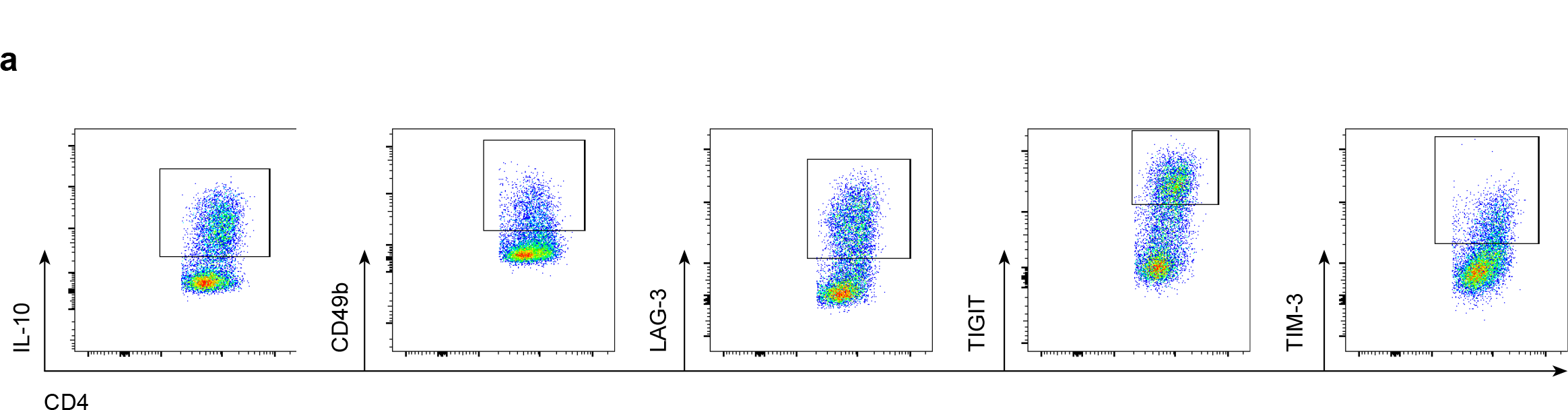
Kidney-derived Foxp3^neg^ IL-10-producing CD4^+^ T cells express CIR **a)** Representative dot plots of IL-10^+^ Foxp3^neg^ cells showing the gating for IL-10^+^, CD49b^+^, LAG-3^+^, TIGIT^+^, and TIM-3^+^ cells corresponding to Figure 4

## Notes

### Competing Interest Statement

The authors have declared no competing interest.

### Summary of Updates

Revised version of the manuscript

## References

1. A. Krebs, C. F., J.-E. Turner, H.-J. Paust, S. Kapffer, T. Koyro, S. Krohn, F. Ufer, M. Friese, R. A. Flavell, B. Stockinger, O. M. Steinmetz, R. A. K. Stahl, S. Huber, and U. Panzer. 2016. Plasticity of Th17 Cells in Autoimmune Kidney Diseases. J. Immunol. 197: 449–457.

2. Tipping, P. G., and S. R. Holdsworth. 2006. T Cells in Crescentic Glomerulonephritis. J. Am. Soc. Nephrol. 17: 1253–1263.

3. Paust, H.-J., J.-E. Turner, J.-H. Riedel, E. Disteldorf, A. Peters, T. Schmidt, C. Krebs, J. Velden, H.-W. Mittrücker, O. M. Steinmetz, R. A. K. Stahl, and U. Panzer. 2012. Chemokines play a critical role in the cross-regulation of Th1 and Th17 immune responses in murine crescentic glomerulonephritis. Kidney Int. 82: 72–83.

4. Steinmetz, O. M., S. A. Summers, P.-Y. Gan, T. Semple, S. R. Holdsworth, and A. R. Kitching. 2011. The Th17-Defining Transcription Factor RORγt Promotes Glomerulonephritis. J. Am. Soc. Nephrol. 22: 472–483.

5. Summers, S. A., O. M. Steinmetz, M. Li, J. Y. Kausman, T. Semple, K. L. Edgtton, D. B. Borza, H. Braley, S. R. Holdsworth, and A. R. Kitching. 2009. Th1 and Th17 cells induce proliferative glomerulonephritis. J. Am. Soc. Nephrol. 20: 2518–2524.

6. Kluger, M. A., M. Luig, C. Wegscheid, B. Goerke, H.-J. Paust, S. R. Brix, I. Yan, H.-W. Mittrücker, B. Hagl, E. D. Renner, G. Tiegs, T. Wiech, R. A. K. Stahl, U. Panzer, and O. M. Steinmetz. 2014. Stat3 Programs Th17-Specific Regulatory T Cells to Control GN. J. Am. Soc. Nephrol. 25: 1291–1302.

7. Paust, H.-J., J.-H. Riedel, C. F. Krebs, J.-E. Turner, S. R. Brix, S. Krohn, J. Velden, T. Wiech, A. Kaffke, A. Peters, S. B. Bennstein, S. Kapffer, C. Meyer- Schwesinger, C. Wegscheid, G. Tiegs, F. Thaiss, H.-W. Mittrücker, O. M. Steinmetz, R. A. K. Stahl, and U. Panzer. 2016. CXCR3+ Regulatory T Cells Control TH1 Responses in Crescentic GN. J. Am. Soc. Nephrol. 27: 1933–1942.

8. Yang, C., X.-R. Huang, E. Fung, H.-F. Liu, and H.-Y. Lan. 2017. The Regulatory T-cell Transcription Factor Foxp3 Protects against Crescentic Glomerulonephritis. Sci. Rep. 7: 1481.

9. Paust, H.-J., A. Ostmann, A. Erhardt, J.-E. Turner, J. Velden, H.-W. Mittrücker, T. Sparwasser, U. Panzer, and G. Tiegs. 2011. Regulatory T cells control the Th1 immune response in murine crescentic glomerulonephritis. Kidney Int. 80: 154–164.

10. Ostmann, A., H. J. Paust, U. Panzer, C. Wegscheid, S. Kapffer, S. Huber, R. A. Flavell, A. Erhardt, and G. Tiegs. 2013. Regulatory T cell-derived IL-10 ameliorates crescentic GN. J. Am. Soc. Nephrol. 24: 930–942.

11. Huber, S., N. Gagliani, E. Esplugues, W. O’Connor, F. J. Huber, A. Chaudhry, M. Kamanaka, Y. Kobayashi, C. J. Booth, A. Y. Rudensky, M. G. Roncarolo, M. Battaglia, and R. A. Flavell. 2011. Th17 Cells Express Interleukin-10 Receptor and Are Controlled by Foxp3- and Foxp3+ Regulatory CD4+ T Cells in an Interleukin-10- Dependent Manner. Immunity 34: 554–565.

12. Groux, H., A. O’Garra, M. Bigler, M. Rouleau, S. Antonenko, J. E. de Vries, and M. G. Roncarolo. 1997. A CD4+T-cell subset inhibits antigen-specific T-cell responses and prevents colitis. Nature 389: 737–742.

13. Gagliani, N., C. F. Magnani, S. Huber, M. E. Gianolini, M. Pala, P. Licona-Limon, B. Guo, D. R. Herbert, A. Bulfone, F. Trentini, C. Di Serio, R. Bacchetta, M. Andreani, L. Brockmann, S. Gregori, R. A. Flavell, and M. G. Roncarolo. 2013. Coexpression of CD49b and LAG-3 identifies human and mouse T regulatory type 1 cells. Nat. Med. 19: 739–746.

14. Brockmann, L., N. Gagliani, B. Steglich, A. D. Giannou, J. Kempski, P. Pelczar, M. Geffken, B. Mfarrej, F. Huber, J. Herkel, Y. Y. Wan, E. Esplugues, M. Battaglia, C. F. Krebs, R. A. Flavell, and S. Huber. 2017. IL-10 Receptor Signaling Is Essential for TR1 Cell Function In Vivo. J. Immunol. 198: 1130–1141.

15. Diefenhardt, P., A. Nosko, M. A. Kluger, J. V. Richter, C. Wegscheid, Y. Kobayashi, G. Tiegs, S. Huber, R. A. Flavell, R. A. K. Stahl, and O. M. Steinmetz. 2018. IL-10 Receptor Signaling Empowers Regulatory T Cells to Control Th17 Responses and Protect from GN. J. Am. Soc. Nephrol. 29: 1825–1837.

16. Roncarolo, M. G., S. Gregori, R. Bacchetta, M. Battaglia, and N. Gagliani. 2018. The Biology of T Regulatory Type 1 Cells and Their Therapeutic Application in Immune-Mediated Diseases. Immunity 49: 1004–1019.

17. Brockmann, L., S. Soukou, B. Steglich, P. Czarnewski, L. Zhao, S. Wende, T. Bedke, C. Ergen, C. Manthey, T. Agalioti, M. Geffken, O. Seiz, S. M. Parigi, C. Sorini, J. Geginat, K. Fujio, T. Jacobs, T. Roesch, J. R. Izbicki, A. W. Lohse, R. A. Flavell, C. Krebs, J. A. Gustafsson, P. Antonson, M. G. Roncarolo, E. J. Villablanca, N. Gagliani, and S. Huber. 2018. Molecular and functional heterogeneity of IL-10- producing CD4 + T cells. Nat. Commun. 9.

18. Pan, L., J. Wang, J. Liu, L. Guo, and S. Yang. 2021. Deficiency in the frequency and function of Tr1 cells in IgAV and the possible role of IL-27. Rheumatol. (United Kingdom) 60: 3432–3442.

19. Tsai, Y. G., J. W. Chien, Y. M. Chiu, T. C. Su, P. F. Chiu, K. H. Hsiao, and C. Y. Lin. 2022. Lupus nephritis with corticosteroid responsiveness: molecular changes of CD46-mediated type 1 regulatory T cells. Pediatr. Res. 92: 1099–1107.

20. Gan, P. Y., D. S. Y. Tan, J. D. Ooi, M. A. Alikhan, A. R. Kitching, and S. R. Holdsworth. 2016. Myeloperoxidase peptide-based nasal tolerance in experimental ANCA-Associated GN. J. Am. Soc. Nephrol. 27: 385–391.

21. Gagliani, N., M. C. Amezcua Vesely, A. Iseppon, L. Brockmann, H. Xu, N. W. Palm, M. R. De Zoete, P. Licona-Limón, R. S. Paiva, T. Ching, C. Weaver, X. Zi, X. Pan, R. Fan, L. X. Garmire, M. J. Cotton, Y. Drier, B. Bernstein, J. Geginat, B. Stockinger, E. Esplugues, S. Huber, and R. A. Flavell. 2015. TH17 cells transdifferentiate into regulatory T cells during resolution of inflammation. Nature 523: 221–225.

22. Hirota, K., J. H. Duarte, M. Veldhoen, E. Hornsby, Y. Li, D. J. Cua, H. Ahlfors, C. Wilhelm, M. Tolaini, U. Menzel, A. Garefalaki, A. J. Potocnik, and B. Stockinger. 2011. Fate mapping of IL-17-producing T cells in inflammatory responses. Nat. Immunol. 12: 255–263.

23. Kamanaka, M., S. T. Kim, Y. Y. Wan, F. S. Sutterwala, M. Lara-Tejero, J. E. Galán, E. Harhaj, and R. A. Flavell. 2006. Expression of Interleukin-10 in Intestinal Lymphocytes Detected by an Interleukin-10 Reporter Knockin tiger Mouse. Immunity 25: 941–952.

24. Esplugues, E., S. Huber, N. Gagliani, A. E. Hauser, T. Town, Y. Y. Wan, W. O’Connor, A. Rongvaux, N. Van Rooijen, A. M. Haberman, Y. Iwakura, V. K. Kuchroo, J. K. Kolls, J. A. Bluestone, K. C. Herold, and R. A. Flavell. 2011. Control of TH17 cells occurs in the small intestine. Nature 475: 514–518.

25. Paust, H.-J., N. Song, D. De Feo, N. Asada, S. Tuzlak, Y. Zhao, J.-H. Riedel, M. Hellmig, A. Sivayoganathan, A. Peters, A. Kaffke, A. Borchers, U. O. Wenzel, O. M. Steinmetz, G. Tiegs, E. Meister, M. Mack, C. Kurts, S. von Vietinghoff, M. T. Lindenmeyer, E. Hoxha, R. A. K. Stahl, T. B. Huber, S. Bonn, C. Meyer- Schwesinger, T. Wiech, J.-E. Turner, B. Becher, C. F. Krebs, and U. Panzer. 2023. CD4 + T cells produce GM-CSF and drive immune-mediated glomerular disease by licensing monocyte-derived cells to produce MMP12 . Sci. Transl. Med. 15: 1–13.

26. Maynard, C. L., L. E. Harrington, K. M. Janowski, J. R. Oliver, C. L. Zindl, A. Y. Rudensky, and C. T. Weaver. 2007. Regulatory T cells expressing interleukin 10 develop from Foxp3+ and Foxp3− precursor cells in the absence of interleukin 10. Nat. Immunol. 8: 931–941.

27. Yogev, N., T. Bedke, Y. Kobayashi, L. Brockmann, D. Lukas, T. Regen, A. L. Croxford, A. Nikolav, N. Hövelmeyer, E. von Stebut, M. Prinz, C. Ubeda, K. J. Maloy, N. Gagliani, R. A. Flavell, A. Waisman, and S. Huber. 2022. CD4+ T-cell-derived IL- 10 promotes CNS inflammation in mice by sustaining effector T cell survival. Cell Rep. 38.

28. Alikhan, M. A., M. Huynh, A. R. Kitching, and J. D. Ooi. 2018. Regulatory T cells in renal disease. *Clin. Transl. Immunol*. 7: e1004.

29. Desreumaux, P., A. Foussat, M. Allez, L. Beaugerie, X. Hébuterne, Y. Bouhnik, M. Nachury, V. Brun, H. Bastian, N. Belmonte, M. Ticchioni, A. Duchange, P. Morel– Mandrino, V. Neveu, N. Clerget–Chossat, M. Forte, and J. Colombel. 2012. Safety and Efficacy of Antigen-Specific Regulatory T-Cell Therapy for Patients With Refractory Crohn’s Disease. Gastroenterology 143: 1207–1217.e2.

30. Bacchetta, R., B. Lucarelli, C. Sartirana, S. Gregori, M. T. Lupo Stanghellini, P. Miqueu, S. Tomiuk, M. Hernandez-Fuentes, M. E. Gianolini, R. Greco, M. Bernardi, E. Zappone, S. Rossini, U. Janssen, A. Ambrosi, M. Salomoni, J. Peccatori, F. Ciceri, and M.-G. Roncarolo. 2014. Immunological Outcome in Haploidentical-HSC Transplanted Patients Treated with IL-10-Anergized Donor T Cells. Front. Immunol. 5.

